# The socioeconomic burden of severe mental disorder in China, 2014-2017: a prevalence study

**DOI:** 10.1101/867432

**Authors:** Mengxue Xie, Ke Ju, Jie Ni, Xiaoheng Zhao, Xiaorong Li, Jay Pan, Qian Zhou

**Author notes:** Correspondence to: Dr Qian Zhou, West China School of Public Health (West China Fourth Hospital), Sichuan University, Chengdu, Sichuan, China.

## Abstract

Mental illness is a chronic disease with high morbidity and mortality rate, resulting in heavy economic burden for family and society, especially the severe ones. Chinese government has taken a series of action on the treatment and management of severe mental disorder (SMD), but the scale of economic burden caused by SMD was still unclear. In this paper, we applied prevalence-based bottom-up approach to estimate the direct and indirect economic burden of SMD in Southwest China from 2014 to 2017. We used the sampled inpatient medical record data of patient with SMD to calculate the total direct cost and estimated the indirect economic burden using human capital approach.The total ecomonic burden of SMD was USD9,733 million and direct burden was contiributed for 7.5% as USD734.5 million in four years total. The growth rate of direct medical cost was declined due to the health policy reform and total cost control policy. The indirect cost was rapidly increased in four years when estimated with DALYs reported in GBD2010 and resulting USD8,998.6 million. We next estimated the indirect burden using sample DALYs and occupation wage cost approach as sensitive analysis. We found that the indirect burden was sensitive to the key variable and the estimation approach, but the estimates share same increasing trend but with different velocity. Our study suggests that SMD in China have posed substantial economic burden at individual and social levels, with the appropriate estimation of total economic burden, our reaserch would attract more attention and be helpful for health resources distribution.

## Background

Mental illness is a chronic disease with high morbidity and mortality rate, resulting in heavy economic burden for family and society [1]. The WHO estimated that mental illness accounted for 10.5% of global burden of diseases in 2011 and would increase to 14.7% in 2020 [2]. The increasing proportion of mental health was basically due to the rapid growth burden in developing country, which was 9.0% in 2011 and 13.7% in 2020[3]. With the increasing prevalence rate, the disease burden of mental illness has become a common concern all over the world, especially for developing countries.

China is the largest developing country in the world, we face serious mental illness challenges [4]. It is reported that “at the end of 2017, over 240 million people in China suffered from all kinds of mental illness and 16 million of which are severe mental disorder” [5]. In the past two decades, Chinese government took a series of action on the treatment and management for severe mental disorder (SMD), such as the “686 Project” starting in 2004 [6] and integrated the services of SMD into primary care in 2009 [7]. Chinese government defined six of the serious mental illness as severe mental disorder (SMD) in 2012 that “severely mental disorder mainly including schizophrenia, schizoaffective disorder, delusional disorder, bipolar disorder, psychotic disorder due to epilepsy and mental retardation” and register patients for long-term management [8]. Since an SMD is a chronic and protracted disease with high recurrence rate, high disability rate and low recovery rate, patients need to repeated consume lots of medical and social health resource for a long time, resulting heavy direct economic burden. At the same time, they need attention and company from their families during the treatment, which lead to the loss of potential healthy labor force. After discharge, due to social discrimination, employment barriers and other reasons, patients' participation in social labor rate is very low, resulting in the lack of dual labor force, resulting in a large number of indirect economic burden.

Previous literatures have discussed the economic costs of some certain types of mental illness in different areas in China, such as depression, dementia and obsession [9, 10, 11], others focused on schizophrenia, one of the most common and serious disease of SMD. It is estimated the direct and indirect economic burden of schizophrenia in Guangzhou Province was RMB 362 million and RMB3.7 billion in total from 2010 to 2012[12]. The total annual costs of mental illness in Shandong Province was increased from RMB10 billion in 2005 to RMB31 billion in 2013, which accounted for 0.7% of total GDP [13]. However, the study of disease burden of SMD in China is insufficient, despite the heavy socioeconomic burden SMD might cause.

Here, we applied the inpatient medical records attributable to SMD first to calculate the direct costs when patients accessed the mental illness serveries, and then estimated indirect economic burden due to productivity losses in Southwest China.

## Methods

### Sources of data

The data were derived from the first page data of inpatient medical record in F Province in Southwest China. We selected patients that were diagnosed with SMD namely schizophrenia, schizoaffective disorder, delusional disorder, bipolar disorder, psychotic disorder due to epilepsy and mental retardation. The first page of medical record documented data on demographics, details of medical costs, disease related information such as ICD-10 diagnosed code and disease severity, and hospital information for all patients. We sampled 5% of inpatient data in each year with the total inpatient data contains 346,171 sample in four years from 2014 to 2017.

Due to the repeatability and long-lasting treatment of SMD, we only included patients who were hospitalized for less than 365 days. Unfortunately, we lost the identification information thus we could not trace down the repeated hospitalization of a SMD patient, so we reported the inpatient costs per person admission in our estimation.

In total, we utilized stratified random sampling method and obtained 18,860 patients who were diagnosed with one of the SMD in 21 cities in F Province.

### Estimation of economic burden

The total economic burden was divided into direct and indirect part, and direct burden contained medical-related costs and non-medical costs [14]. We applied the bottom-up cost calculation method to calculate the direct economic burden: extrapolated the total inpatient costs from sampling costs [15, 16]. Direct medical costs refer to the utilization of outpatient and inpatient services, such as medication in outpatient, treatment, diagnosis and rehabilitation in admission. Direct non-medical costs are corresponded to informal care, which mean the daily expenses during hospitalization for both the patient and his/her families. We calculated the sum of the inpatient expenses such as treatment fee, medication fee and diagnose fee as direct medical costs, then estimated the costs for total SMD inpatients from 2014 to 2017. Due to the limitation of our sample, we could not obtain the outpatient information since the first page of medical record data only track the patients in admission. We used the open database issued by Chinse Government to estimate the total outpatient number and costs. We omit the direct non-medical costs due to the sample information.

As for indirect burden, we estimated the loss of labor productivity due to disease-related disability and mortality with human capital approach [17]. The disability-adjusted life year (DALYs) is a comprehensive and effective index to estimate indirect burden of disease, and has been applied in the study of GBD throughout the world [18, 19].

We calculated the DALYs of SMD using 7.4% that reported in GBD 2010[20]. The age productivity of SMD was calculated based on the age-specific distribution reported in Fifth National Census 2000 and the total admissions [21]. Gross domestic product (GDP) and other relevant statistics were obtained from China Statistical Yearbook from 2015 to 2018. The monetary unit in our paper has been converted to US dollar based on the exchange rate of Bank of China in January 2019 which was 1 dollar to 6.8 RMB.

### Sensitivity Analysis

We carried out two types of sensitivity analysis on the indirect economic burden to test the robustness of our estimation. First, we used the formula described in GDB 2010 to estimate the DALYs of SMD by our sample with native data instead of the average estimation of mental illness DALYs in GBD2010[20]. Next, we used the patients’ occupation wage to estimate the productivity loss for each admission instead of using age productivity weight and per capita GDP. The average daily income was derived from were derived from F Province Statistical Yearbook 2015-2018^1^.

## Results

### Summary statistics of the study sample

It is shown in Table 1 that, of the 17,918 patients with SMD, the mean age was 43.8 years, 3.3% were aged below 18 years, 75.3% were aged between 18–55 years, 21.4% were aged above 55 years. 63.2% were men, and 62.1% were rural residents. Schizophrenia (74.2%) was the most common SMD among the sample, followed by mental retardation (16.2 %), bipolar disorder (7.8%), and other severe mental disorders (1.9%). The total enrollment rate of health insurance was 92%. The proportion of SMD inpatient was increased from 14.1% in 2014 to 32% in 2017, based on the sample method, we inferred that the total inpatient number was increased year by year.

**Table 1.**
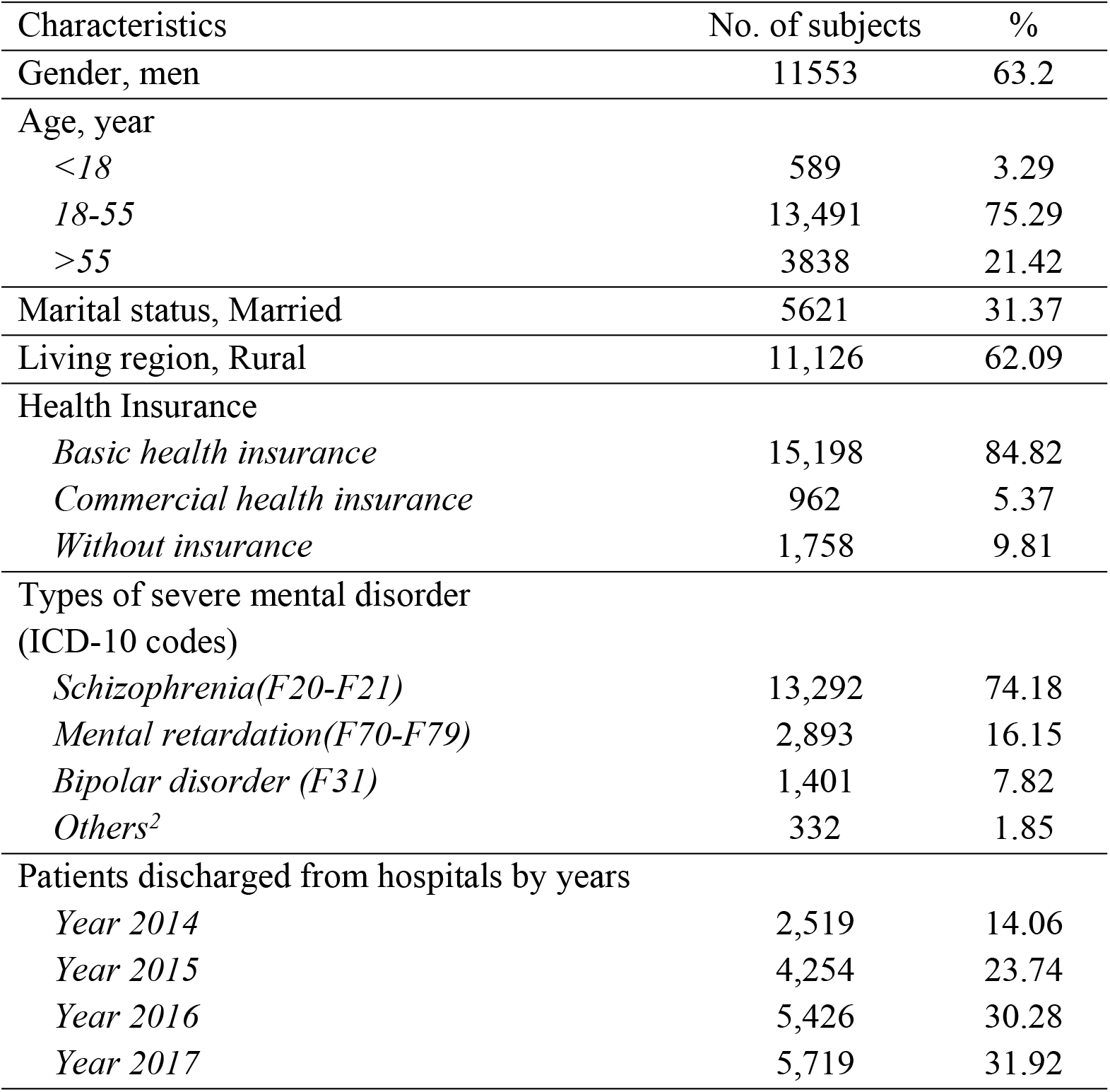
Characteristics of the study sample (n=17,918)

Total annual costs per admission decreased from $2308.8 in 2014 to $1837.2 in 2017. The treatment cost accounts for almost half of total medical cost, but had decreased year by year. The medication cost had decreased from $197.2 in 2014 to $104.8 in 2017, but the examination cost had increased from $235.6 in 2014 to $298.8 in 2017.As for the OOP cost, the average number was decreased to $1100.4 in 2016 than previous years, but increased slightly to $1266.1 in 2017. Generally speaking, the direct economic burden to the patient with SMD had relieved in the past four years.

Young and middle-aged patients accounted for 75.7% of the total costs, specifically 29.2% for those 18 to 39 years old, and 46.5% for those 39 to 55 years old. But the average cost for patients aged below 55 years was decreased year by year, while that of the patients aged above 55 years old was increased.

Patients with schizophrenia accounted for 40.5% of the total costs, 29.7%, 6.8% and 2.4% for that with mental retardation, bipolar disorder, and other SMD. The overall trend of the average cost for patients with severe mental disorders was declined, but with different rate: bipolar disorder and schizophrenia declined 7.8% and 13% in 2017 compared to 2014, mental retardation and others declined 43.1% and 22.5%.

**Table 2.** Medical cost to SMD per patient with different characteristics

| Years | Costs ratio | 2014 | 2015 | 2016 | 2017 |
| --- | --- | --- | --- | --- | --- |
| Inpatient cost |  | 2308.8 | 2064.1 | 1971.2 | 1837.2 |
| *Treatment cost* |  | 1343.4 | 1032.8 | 852.7 | 861.8 |
| *Examination cost* |  | 234.6 | 267.3 | 275.9 | 298.8 |
| *Medication cost* |  | 197.2 | 146.2 | 130.7 | 104.8 |
| Out-of-Pocket cost |  | 1585.9 | 1371.8 | 1100.4 | 1266.1 |

<sup>2</sup> Others included epileptic psychosis, schizoaffective disorder and paranoid psychosis

|  |  |  |  |  |  |
| --- | --- | --- | --- | --- | --- |
| Gender |  |  |  |  |  |
| Male |  | 2473.4 | 2153.4 | 2094.1 | 1979.6 |
| Female |  | 1998.4 | 1912.7 | 1770.7 | 1587.5 |
| Age |  |  |  |  |  |
| Below 18 years old |  | 2033.3 | 1929.6 | 1454.3 | 1367.8 |
| 18 to 55 years old | 75.7% | 2431.6 | 2130.8 | 1910.0 | 1780.9 |
| Above 55 years old |  | 1967.7 | 1845.0 | 2273.5 | 2089.7 |
| Living area |  |  |  |  |  |
| Urban |  | 1699.5 | 1662.3 | 1698.0 | 1606.8 |
| Rural |  | 2635.4 | 2345.0 | 2167.7 | 1951.2 |
| Mental illness |  |  |  |  |  |
| Schizophrenia | 40.5% | 2403.7 | 2060.7 | 2087.8 | 2092.0 |
| Mental retardation | 29.7% | 2285.5 | 2191.6 | 1626.2 | 1299.9 |
| Bipolar disorder | 6.8% | 1507.6 | 1490.3 | 1467.6 | 1390.3 |
| Others | 2.4% | 2325.2 | 2237.8 | 2196.3 | 1802.5 |

The average cost for female patient was less than USD2,000 during these four years, and male patient costed over USD2,000. Medical costs were higher for rural patients than urban patients, which means the direct economic burden for male and rural patients are heavier. But it is also shown that the direct economic burden was generally decreased for both male and female, urban and rural patients. As shown in Fig.2, rural patients accounted for over 50 percent of the average cost and urban patients accounted less than 50 percent. But the proportion of rural patients was declined during the four years while urban patients increased, which means the gap between rural and urban narrowed year by year, along with the declining of the average medical cost per patient.

**Fig. 1.**
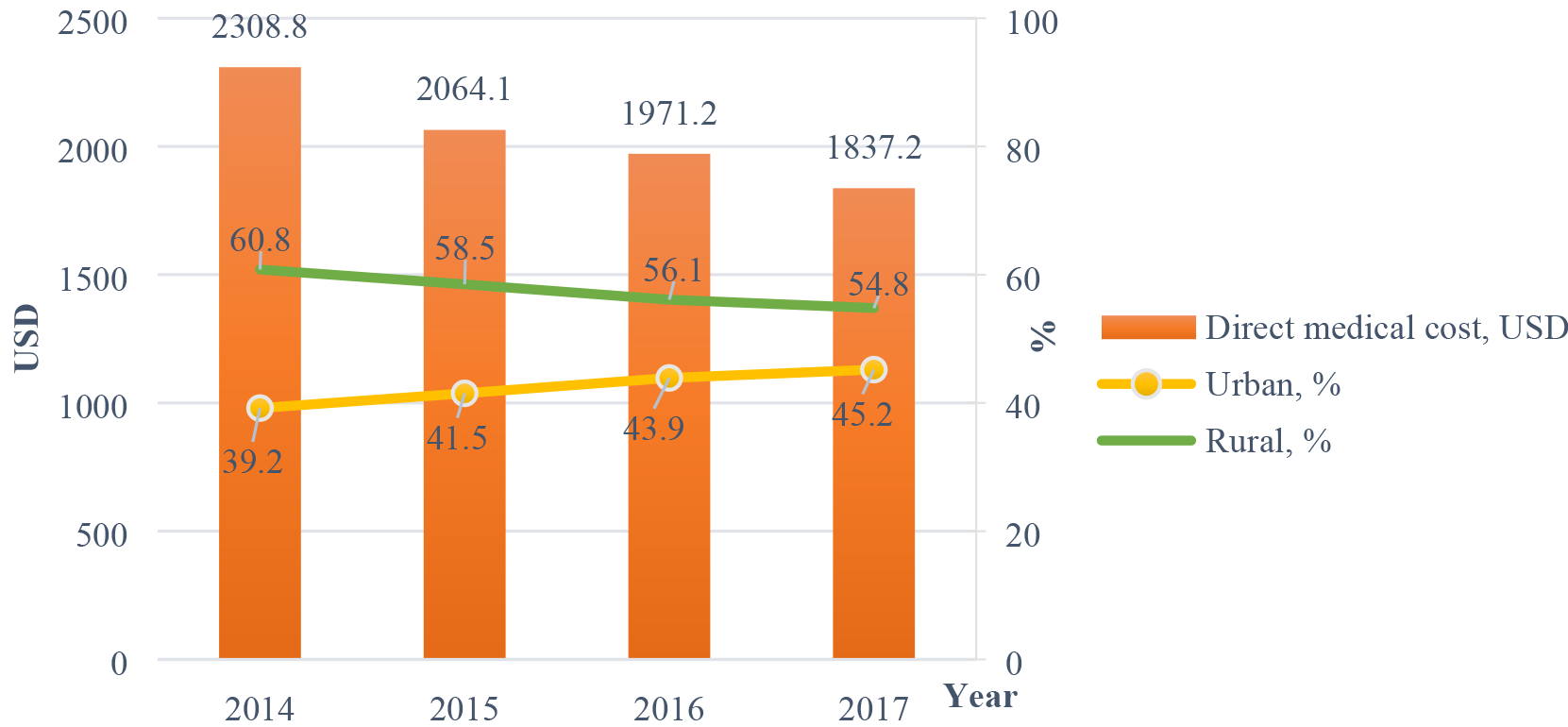
The actual cost of SMD in China. The average inpatient medical cost of SMD(bars, in $) and cost proportion of urban and rural patients over the total cost(lines, %)

**Fig. 2.**
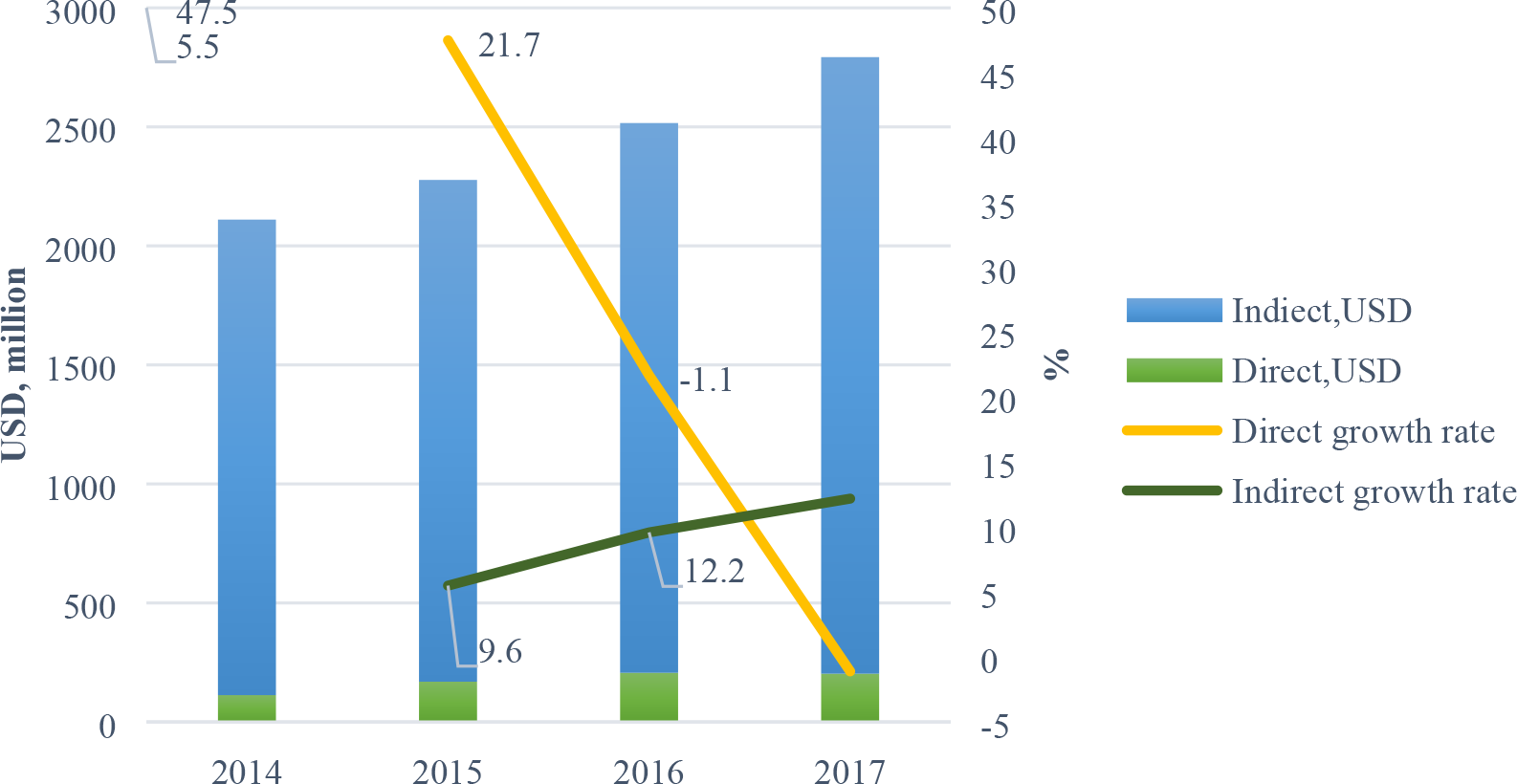
The estimated economic burden of SMD in China. The economic burden of SMD(bars, in $) and variation trend over last year(lines, %)

### Direct economic burden

We divided the estimation of direct economic burden into two aspects. First, we estimated the total inpatient economic burden for patients with SMD during 2014 to 2017 with the sampling data. The total number of inpatients was 346,171 and the total inpatient cost was USD 691.7million (Table 3). The direct inpatient economic burden of each year significantly increased from $113 million in 2014 to $203 million in 2017 while the average medical costs decreased. The direct cause for the increased total economic burden was the rapid growth of the inpatient number which had tripled during the four years.

**Table 3.** Total direct economic burden for patients with SMD in 2014-2017 (USD)

| Items | In total | 2014 | 2015 | 2016 | 2017 |
| --- | --- | --- | --- | --- | --- |
| Total direct costs, \$ in million | **734.5** | **121.6** | **179.3** | **218.2** | **215.6** |
| Numbers of inpatient | 346,171 | 48,810 | 82,403 | 104,890 | 110,428 |
| Inpatient cost, \$ in million | **691.7** | **112.7** | **169.3** | **206.8** | **202.9** |
| *Treatment cost* | 335.3 | 65.6 | 85.1 | 89.4 | 95.2 |
| *Examination cost* | 95.4 | 11.5 | 22.0 | 28.9 | 33.0 |
| *Medication cost* | 47.0 | 9.6 | 12.0 | 13.7 | 11.6 |
| Numbers of outpatient, in million<sup>a</sup> | 1289.5 | 295.6 | 304.2 | 330.9 | 358.8 |
| Average outpatient cost<sup>b</sup> | 32.7 | 29.6 | 32.2 | 33.9 | 34.9 |
| Outpatient cost, \$ in million | **42.8** | **8.9** | **10.0** | **11.4** | **12.7** |

Note a, b: Data source: China Health Statistics Yearbook 2015 to 2018, at www.nhfpc.gov.cn

Then we estimated the total outpatient costs using China Health Statistics Yearbook 2015 to 2018 data. The total number of outpatients was USD 1289.5 million during these years, and the average outpatient cost was increased from USD 32.7 to USD 34.9. Note that the outpatient cost of SMD was not reported in the statistic yearbook, we used the average outpatient cost of whole disease as a proxy. The outpatient cost was estimated to be USD 42.8 million in total, and with 10% growth rate in each year.

At last, we simply combined the inpatient and outpatient cost to obtain the total direct economic burden, which was USD 734.5 million in total and 121.6 million, 179.3 million, 218.2 million and 215.6 million respectively.

### Indirect economic burden

The indirect economic burden within these four years was 8.9 billion in total, with average 7.4% DALYs and 0.641 age productivity weight^3^ in Table 5. The indirect economics burden increased from USD1,998 million to USD2,590 million with 10% growth rate each year due to the growing population and per capita GDP, while the age productivity weight has decreased slightly. In fact, the DALYs will not stay unchanged during these four years, thus we estimated the DALYs of SMD in each year using our sample in the sensitivity analysis later.

**Table 5.** Estimation of indirect economic burden of patients with SMD

| Items | Total | 2014 | 2015 | 2016 | 2017 |
| --- | --- | --- | --- | --- | --- |
| Total indirect cost, \$ in million | **8,998.6** | 1,997.7 | 2,107.8 | 2,309.1 | 2,589.9 |
| Population | 329,080 | 81,400 | 82,040 | 82,620 | 83,020 |
| Per Capita GDP<sup>a</sup> | 5,755.8 | 5,165.9 | 5,408.1 | 5,882.8 | 6,566.4 |
| DALYs<sup>b</sup> | 7.4 | 7.4 | 7.4 | 7.4 | 7.4 |
| Age productivity weight<sup>c</sup> | 0.641 | 0.662 | 0.646 | 0.64 | 0.641 |

Note: a: data source: China Statistical Yearbook at www.stats.gov.cn
b: DALYs was estimated in GBD2010
c: estimation provided in Appendix Table 1

### Total economic burden

The total economic burden of SMD in Southwest China was estimated to be USD 9.7 billion during 2014 to 2017 in Table 6. Direct economic burden only contributed 7.5% of total economic burden while the productivity losses accounted the remaining 92%. The total economic burden increased from 2.1 billion in 2014 to 2.8 billion in 2017, with the average growth rate 9.8%. We noticed that the growth rate of direct cost was decreased from 47.5% to 21.5% and started to negative growth in 2017(Fig.2). The relived economic burden for SMD patients is inseparable from the efforts of the government.

**Table 6.** Estimated economic burden of SMD in China, 2014-2017

| Items | Total | 2014 | 2015 | 2016 | 2017 |
| --- | --- | --- | --- | --- | --- |
| Direct cost, \$ in million | 734.5 | 121.6 | 179.3 | 218.2 | 215.6 |
| Inpatient cost | 691.7 | 112.7 | 169.3 | 206.8 | 202.9 |
| Outpatient cost | 42.8 | 8.9 | 10.0 | 11.4 | 12.7 |
| Indirect cost, \$ in million | | | | | |
| DALYs cost | 8,998.6 | 1,997.7 | 2,107.8 | 2,309.1 | 2,589.8 |
| Total economic costs, \$ in million | **9,733.10** | 2,119.30 | 2,287.10 | 2,527.30 | 2,805.40 |

On the other hand, the growth rate of indirect economic burden was increased, from 5.5% in 2015 to 12.2% in 2107, compared to last year. The estimation of indirect growth rate was based on per capita GDP and the total inpatient number, a rapid growth of these two factors resulting the increasing of indirect economic burden. Thus, the increasing of major index in the estimation leading the indirect economic burden of SMD upwards, and this trend will not stop as long as the economic and social development continues.

### Sensitivity analysis

The results of sensitivity analysis were revealed in Table 7. We found that the estimated indirect burden would change significantly with the change of key index. In the early estimation, we used the worldwide average DALYs reported in GBD2010, which may had underestimated the effect of SMD in China, especially in Southwest China since the prevalence is higher in underdeveloped areas [22]. In that case, first we calculated the sample DALYs as 17.6% in F province and found that the indirect economic burden escalated from USD 9.7 billion to USD 30.4 billion.

**Table 7.** Sensitivity Analysis on economic burden of SMD

| Cost Items | Total | 2014 | 2015 | 2016 | 2017 |
| --- | --- | --- | --- | --- | --- |
| Indirect cost, \$ in million | | | | | |
| DALYs cost | 8,998.6 | 1,997.7 | 2,107.8 | 2,309.1 | 2,589.8 |
| Sample estimation DALYs cost | 21,368.6 | 4,899.4 | 5,044.5 | 5,475.7 | 6,150.0 |
| Occupation wage cost | 48,097.7 | 5,575.5 | 10,633.1 | 15,401.7 | 19,055.8 |

Next, we used alternative method to estimate productivity losses by the occupation of the patient. We summarized the occupations into three categories: employee, low-income and unemployment. Employee means patient was employed by an employer, and low income contained patients that were farmers and self-employments. The wage information of employee and farmer was reported in the China Statistical Yearbook 2015 to 2018. For unemployment patient that were unemployed, students and retired patients, we used the monthly minimum wage as the proxy index for occupational wage. We find that the indirect economic burden boosted from previous estimates USD 9 billion and USD 21 billion into 48 billion, which was approximately 2.54% of the total GDP in four years of F province.

## Discussion

Our estimation of the total economic burden of SMD increased from USD2.1 billion in 2014 to USD2.8 billion in 2017, with average 9.8% growth rate. The direct economic burden increased from USD121.6 million in 2014 to USD218.2 million in 2016 while decreased slightly to USD215.6 million in 2017 due to the SMD management system reform and medical expenses control policy [23,24,25]. The SMD management system reform released potential demand for mental health services and medical expenses control policy managed hospital behaviors, resulted the increasing admissions and decreasing per patient costs.

The annual indirect economic costs of SMD estimated with reported DALYs within these four years was USD2.3 billion, was equivalent to 0.5% of the average GDP. Meanwhile, the annual indirect economic costs estimated by using sample DALYs and occupation wage costs were 1.1% and 2.7% of the average GDP, which suggested that we underestimated the indirect economic burden in our initial estimation. In Europe, costs of mental disorders were estimated to account for 3.5 % of GDP [28]. The total economic burden of mental health in Pakistan to be PKR 165.9 billion (USD 2.8 billion) in 2005-06, which was 2.5% of the gross domestic product (GDP) of Pakistan [29].

Compared with some foreign studies [28, 29, 30], our estimation is lower, but it is much higher than the research results of domestic literature. First of all, most of the data used in the domestic literatures are from prefecture-level cities or schizophrenia inpatients in local hospitals, so the sample size of admission is small and there is a certain bias [31, 32]. In our study, for the first time, the Provincial sampling data is used to estimate the provincial data according to the sampling proportion, which improves the accuracy of sample number and greatly reduces the estimation error of economic burden. Even though some [33] used mortality rate in Yunnan Province to estimate the economic burden of schizophrenia which was nearly RMB4.6 billion in Southwest China, the results would lead to great bias due to the data accuracy and the uniqueness of one SMD. It is reasonable to believe that our estimation is more accurate than the existing research in China.

The indirect economic burden of SMD was greater than the direct economic burden. First of all, most of SMD in our sample were aged from 14 to 59, more males than females, which lead to 0.64 of the age productivity weight and brings serious losses to social production. Secondly, SMD is easy to recur and difficult to cure. After discharge, it is accompanied with low labor force, low employment rate and life expectancy. According to an estimation of 17 kinds of common severe diseases in Beijing [34], cerebral vascular diseases have the highest DALYs, which is 15.77-person years per thousand people. The estimated SMD DALYs in 2014-2017 in our sapmple is 17.64-person years per thousand people, which is higher than that of common severe diseases. It can be seen that schizophrenia brings serious life loss to the residents of our province, which deserves more attention and research.

Our study still has several limitations though. First, considering that there existed great differences in economic development, population structure and medical service capacity across the country, it is insufficient to estimate the economic burden of SMD only in one province in China. However, the fact that mental illness especially SMD is more seriously in Southwest China, which makes our study valuable to draw more attention to these less-developed regions. Second, we might underestimate the total economic burden because we were not able to account for certain costs due to lack of related information. As for the direct medical costs, we calculated the total inpatient costs directly but estimated the total outpatient costs with open database issued by Chinese Government. It is an effective and reliable method to fulfill the estimation of direct economic burden, but we still hope the actual outpatient data of SMD would be valid in the near future so that we can calculate the truly total direct economic burden. We treated the direct non-medical costs as zero because the data did not provide the information related to informal care and accidental and non-accidental injuries costs attributable to SMD, which could definitely lead a downward bias to our estimation. Third, the estimation of indirect burden lack of robustness. The estimated indirect burden boosted from 9 billion to 43 billion when we using GBD2010 reported DALYs, sample DALYs and occupational wage for a sensitivity test in Table 7, which suggested the key variable in the estimation should be carefully chosen.

## Conclusion

Our study suggests that SMD in China have brought huge economic burden for family and society, and direct medical cost contributed for 7.5% of total economic cost with the ratio to indirect costs 1:12.3. The proportion of direct and indirect burden suggested that the input to the medical services for severe mental disorder was insufficient, compared to the direct and indirect ratio of 20% in existing researches[‥]. Besides, the social economic impact would be much greater when the prevalence of SMD and per capita GDP increased. Although our estimate is not accurate, we made a valuable attempt to estimate the disease burden of SMD, in particularly the Chinese government issued new management regulations to strengthen the management of SMD in 2018[36]. With the appropriate estimation of total economic burden, studies like ours can attract more researches and be helpful for health resources distribution.

## Author Contributions

All authors were responsible for the structure of this paper. MX and QZ conceived and designed the study. MX and QZ carried out the analysis and wrote the paper. XZ, JN, XL, and JP handled the data and sought funding. All authors have seen and approved the final version of submission.

## Appendix

**Appendix Table 1.** Age productivity weight of patients with SMD

|  | Numbers of sample | Age productivity weight of patients with SMD | 0~14 | 15~49 | 50~59 | 60~ |
| --- | --- | --- | --- | --- | --- | --- |
| Total Four years | 17918 | **0.641** | 327 | 11601 | 3135 | 2855 |
| Year 2014 | 2519 | **0.662** | 57 | 1607 | 524 | 430 |
| Year 2015 | 4254 | **0.646** | 90 | 2799 | 733 | 632 |
| Year 2016 | 5426 | **0.640** | 91 | 3624 | 836 | 848 |
| Year 2017 | 5719 | **0.641** | 89 | 3571 | 1114 | 945 |

Note: The age productivity weight was 0 for aged 0~14, 0.75 for aged 15~49, 0.8 for aged 50~59, 0.1 for aged over 60, according to the China Fifth National Census, 2000.

**Appendix Table 2.**
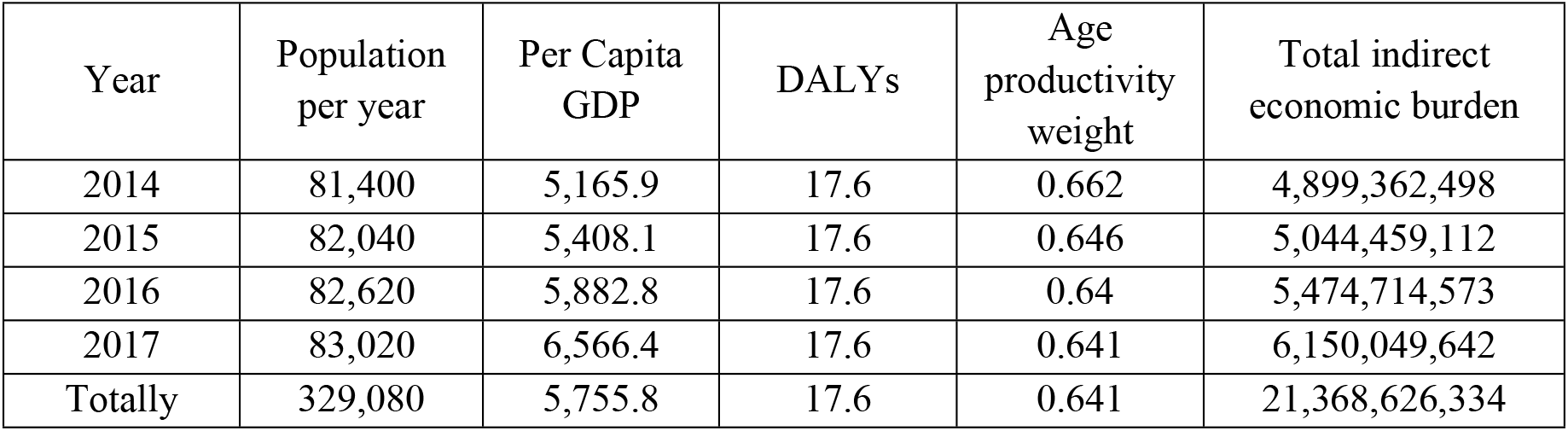
Estimation of indirect economic burden using sample DALYs

| Year | Population per year | Per Capita GDP | DALYs | Age productivity weight | Total indirect economic burden |
| --- | --- | --- | --- | --- | --- |
| 2014 | 81,400 | 5,165.9 | 17.6 | 0.662 | 4,899,362,498 |
| 2015 | 82,040 | 5,408.1 | 17.6 | 0.646 | 5,044,459,112 |
| 2016 | 82,620 | 5,882.8 | 17.6 | 0.64 | 5,474,714,573 |
| 2017 | 83,020 | 6,566.4 | 17.6 | 0.641 | 6,150,049,642 |
| Totally | 329,080 | 5,755.8 | 17.6 | 0.641 | 21,368,626,334 |

**Appendix Table 3.** Average monthly wage in F province in from 2014 to 2017.

|  | total | 2014 | 2015 | 2016 | 2017 |
| --- | --- | --- | --- | --- | --- |
| employee | 5094.1 | 4354.6 | 4909.6 | 5327.2 | 5784.9 |
| farmer | 4147.1 | 3279.8 | 3977.4 | 4401.1 | 4929.9 |
| unemployment | 1475.0 | 1400 | 1500 | 1500 | 1500 |

**Appendix Table 4.** Total cost of SMD attributed to occupation wage

|  | Average daily wage | Average admission days | # of inpatient | Cost |
| --- | --- | --- | --- | --- |
| employee | 25.0 | 80.0 | 16035 | 32,070,000.0 |
| low-income | 20.3 | 85.0 | 225770 | 389,566,135.0 |
| unemployment | 7.2 | 79 | 104327 | 59,341,197.6 |

## Footnotes

1 Data source: http://tjj.sc.gov.cn/

2 Others included epileptic psychosis, schizoaffective disorder and paranoid psychosis

3 We show our estimation of age productivity weight of SMD in Appendix 1.

